# Down regulation of the liver lipid metabolism induced by hypothyroidism in mice: metabolic flexibility favors compensatory mechanisms in white adipose tissue

**DOI:** 10.1101/2022.12.22.521639

**Authors:** Lamis Chamas, Isabelle Seugnet, Odessa Tanvé, Valérie Enderlin, Marie-Stéphanie Clerget-Froidevaux

**Author notes:** These authors jointly supervised this work (Valérie Enderlin, Marie-Stéphanie Clerget-Froidevaux).

## Abstract

In mammals, the maintenance of energy homeostasis relies on complex mechanisms requiring tight synchronization between peripheral organs and the brain. Thyroid hormones (TH), among their pleiotropic actions, play a central role in these regulations. Hypothyroidism, which is characterized by low circulating TH levels, slows down the metabolism, leading to a reduction in energy expenditure, as well as in lipid and glucose metabolism, and to insulin resistance. Our objective was to evaluate whether metabolic deregulations induced by hypothyroidism could be avoid by regulatory mechanisms involved in metabolic flexibility. To this aim, we compared the response to hypothyroidism in two mouse strains, the wild-derived WSB/EiJ mouse strain characterized by a diet-induced obesity (DIO) resistance due to its high metabolic flexibility phenotype and the C57BL/6J mice, prone to DIO. Adult mice were fed with a low-iodine diet supplemented with 6-n-propyl-2-thiouracyl (PTU) for 7 weeks to induce hypothyroidism. Our results show that hypothyroidism, characterized by a decrease in serum T4 levels, led to metabolic deregulations, as an alteration of lipid metabolism in the liver of both strains. However, the decrease in hepatic lipid synthesis was compensated in WSB/EiJ mice by a mobilization of lipid reserves from white adipose tissue, but not in the C57BL/6J mice. No peripheral or hypothalamic inflammatory response to hypothyroidism was observed in both strains. Moreover, gene expression analysis showed that hypothyroidism stimulates the hypothalamic orexigenic circuit in both strains, but unchanged Mc4r and LepR expression in hypothyroid WSB/EiJ mice strain, which reflect their adaptability to maintain their body weight, contrary to C57BL/6J mice. Our results show that WSB/EiJ mice displayed a phenotype of resistance to metabolic dysregulations induced by hypothyroidism, by compensatory mechanisms. This response as well as their resistance to HFD-induced obesity highlights their adaptive capacities to maintain metabolic homeostasis, namely, their high metabolic flexibility, despite serum hypothyroidism. This model sheds light on the importance of local thyroid homeostasis to maintain lipid metabolism and metabolic homeostasis.

## Introduction

In mammals, the maintenance of energy homeostasis relies on complex mechanisms requiring tight synchronization between peripheral organs and the brain. It involves both the endocrine and central nervous system (CNS). The hypothalamus plays an essential role in the central control of energy balance, especially as a nutrient sensor modulating food intake and energy expenditure accordingly to metabolic status (review in (Flier, 2004)). It orchestrates a dynamic crosstalk between its different nuclei and the peripheral metabolic tissues. The first order neurons present in the arcuate nucleus (ARC) integrate peripheral signals such as hormones (leptin, insulin, etc.) and nutrients (lipids, glucose) in order to monitor metabolic status of the entire organism (review in (Flier, 2004; Schwartz, 2006)). In response, potent outputs are conveyed through orexigenic (NPY/AGRP) or anorexigenic (POMC/CART) neurons to the second order neurons located in other nuclei of the hypothalamus as the paraventricular nucleus (PVN), which will thus restore energy homeostasis, ensuring weight maintenance by modulating energy expenditure and food intake (Le Thuc *et al*., 2017). One of the main actors in maintaining this energy balance is the hypothalamo-pituitary-thyroid axis. This axis controls the synthesis of the thyroid hormones (THs), which, among their pleiotropic actions, are key regulators of the energy balance (review in (Mullur, Liu and Brent, 2014)). Thyroxine (T4) and the transcriptionally active triiodothyronine (T3) ensure metabolic homeostasis of the whole body mainly through the transcriptional activity of their nuclear receptors. They control all aspects of metabolism, acting both centrally and in peripheral metabolic organs (particularly the liver and adipose tissues) (Cicatiello, Di Girolamo and Dentice, 2018). THs act on lipid and carbohydrate metabolism, as well as on metabolic-cellular mechanisms such as mitochondrial activity and thermogenesis, and also on food intake and energy expenditure (Song, Yao and Ying, 2011; Mullur, Liu and Brent, 2014). Due to the high implication of THs in metabolic control, any disruption of TH homeostasis could be a cause of metabolic disorders. Hypothyroidism, which is characterized by low circulating TH levels, slows down the metabolism, leading to a reduction in energy expenditure, as well as in lipid and glucose metabolism, and to insulin resistance (Brenta, 2011; Mullur et al., 2014; Mcaninch & Bianco, 2014). However, other factors are able to disrupt energy homeostasis, as food rich in fat and sugar, which also lead to metabolic dysregulations and the development of metabolic disorders, the most common being obesity and insulin resistance (Liu et al., 2015; De Souza et al., 2005; review in (Guillemot-legris and Muccioli, 2017)). Research on these metabolic disorders commonly uses the diet induced obesity (DIO) model, which consists in feeding mice a high fat diet (HFD) in order to induce obesity and metabolic syndrome. Interestingly, some mouse strains are resistant to DIO, as the WSB/Eij strain, which could thus be a very useful model to unravel pathways involved in metabolic homeostasis. Indeed, these mice depict an extraordinary ability to maintain metabolic homeostasis even under HFD (Lee *et al*., 2011; Terrien *et al*., 2019), while paradoxically exhibiting low circulating levels of T4 (around 3 μg/dL), even still in the reference range obtained from 28 mouse strains (ranging from 2.61 to μg/dL in male mice (Li *et al*., 2008)). This constitutes an interesting enigmatic paradox making WSB/EiJ mice a relevant model for metabolic studies. Thus, regarding the role plays by HT in the control of metabolism, in this article, we were interested in evaluating if WSB/EIJ mice were also able to adapt to the metabolic unbalance induced by hypothyroidism, in comparison with C57BL/6J mice.

For this purpose, we induced a transient period of hypothyroidism in adult C57BL/6J and WSB/EiJ male mice by administrating a commonly used antithyroid molecule, the propylthiouracil (PTU) for seven weeks (Decherf *et al*., 2010; Chaalal *et al*., 2014). We explored the consequences of PTU-induced hypothyroidism on peripheral and central metabolism, by following weight changes, markers of lipid metabolism, as well as expression of genes involved in metabolic pathways in peripheral tissues and in the hypothalamus. As metabolic unbalance is known to induce hypothalamic inflammation (Thaler *et al*., 2012; Terrien *et al*., 2019), we also verified the inflammatory status of the hypothalamus by immunohistochemistry. We showed first that 7 weeks of PTU induced alteration in lipid metabolism, particularly a reduction in hepatic lipid synthesis in both strains. Furthermore, contrary to the C57BL/6J mice, the WSB/EiJ mice were resistant to the metabolic dysregulations induced by hypothyroidism, mainly through an enhanced lipid metabolism in adipose tissue.

## Results

### I. Evaluation of thyroid status of mice after PTU treatment

We first evaluated thyroid status between the two mouse strains after 7 weeks of PTU treatment by measuring circulating T4 levels (Fig.1). Euthyroid WSB/EiJ mice depicted a lower circulating total T4 [3.4 μg/dL] than C57BL/6J mice [6.1 μg/dL] (p= 0.0003). In response to PTU treatment, circulating total T4 levels were decreased by 90% and 80% in C57BL/6J and WSB/EiJ mice [0.06 μg/dL], respectively, compared to their control mice (p < 0.001). These results confirmed that both strains were peripherally hypothyroid after 7-weeks of PTU treatment.

**Figure 1.**
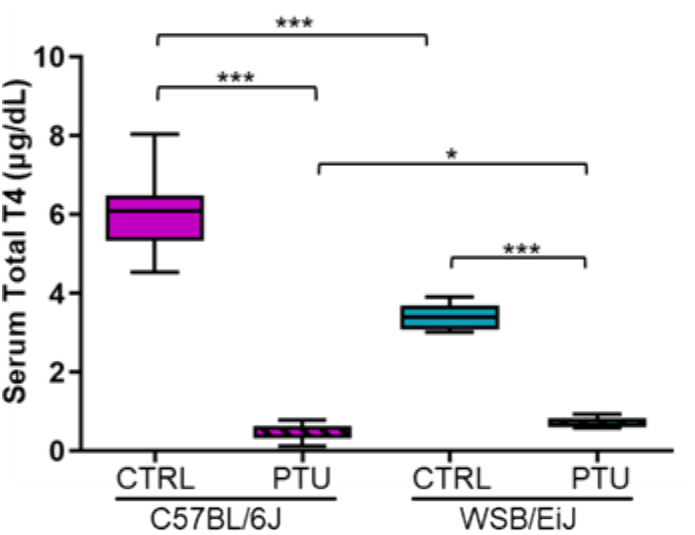
PTU induces hypothyroidism in both mouse strain. Serum total T4 levels were reduced after 7-weeks of PTU treatment in C57BL/6J and WSB/EiJ mice compared to the euthyroid control mice (CTRL) (n = 8-7 per group, non-parametric two-way ANOVA with permutations test). Boxplot represents median values and min-max whiskers. Post Hoc tests results are indicated on the graph (*, p ≤ 0.05; ***, p ≤ 0.001).

### II. Hypothyroidism consequences on metabolism and energy balance

#### 2.1. Body weight, food intake, body fat mass

Once peripheral hypothyroidism was verified, we investigated the consequences of hypothyroidism on peripheral and central metabolism between the two strains. We first evaluated the effect of PTU treatment on the body weight of both mouse strains, particularly the variation of the weight during the treatment (Fig.2A). We first observed a difference in the body weight of euthyroid compared to hypothyroid groups starting from the 1^st^ week of treatment (0.0001 ≤ p ≤ 0.001). The body weight of euthyroid C57BL/6J and WSB/EiJ mice significantly increased in a similar way throughout the treatment, reflecting the growth period of young adult mice (0.0001 ≤ p ≤ 0.05). However, this weight gain was inhibited by PTU treatment as body weight of hypothyroid C57BL/6J mice was significantly decreased throughout the weeks and the diminution reached 6% of the initial weight at the 5^th^ week of treatment (0.0001 ≤ p ≤ 0.05). In contrast, the body weight of hypothyroid WSB/EiJ mice was maintained throughout the treatment (p > 0.05), although their body weight was lower than their control group’s weight, which increased over the weeks (0.0001 ≤ p ≤ 0.001). The PTU treatment induced different effect between the hypothyroid groups: C57BL/6J mice lost weight whereas WSB/EiJ maintained their body weight during the treatment.

**Figure 2.**
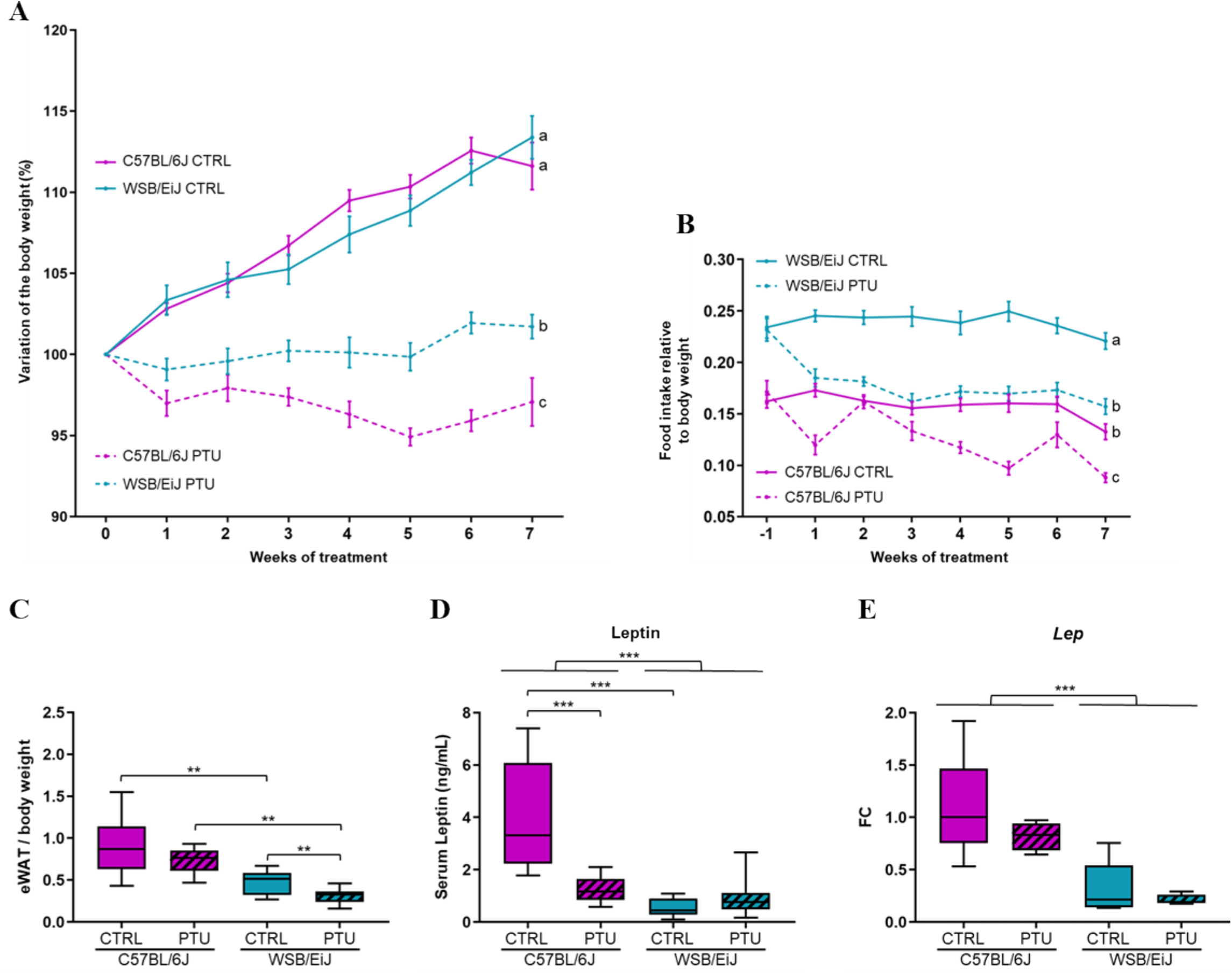
PTU-treatment effects on metabolism including body weight, food intake and body fat mass. **(A)** Variation upon weeks of treatment of the body weight expressed as percentage of the starting (Week 0) body weight for each group (% each week BW/ Initial BW before the treatment; n =13-14 per group). **(B)** Food intake measurements (relative to BW) during the 7-weeks of treatment (n = 13-15 per group). Food intake data were collected starting from 24h before the treatment (−1). For (A-B) statistical differences between treatment and time were assessed by two-way repeated measures ANOVA followed by Tukey’s multiple-comparisons test. Significant differences were indicated by different letters to account for group differences at week 7. Values are mean ± SEM. **(C)** Epididymal white adipose tissue (eWAT) weights measured at the end of the treatment (relative to final BW; n = 12-15 per group; non-parametric one-way ANOVA with permutations test). **(D)** Circulating leptin (in ng/mL; n = 9-13 per group) and **(E)** its mRNA expression levels in eWAT (n = 5-6 per group) evaluated after 7-weeks of treatment. Data are represented as relative fold-change expression (FC). Boxplot represents median values and min-max whiskers. Non-parametric two-way ANOVA with permutations test were performed for (D-E) data (see table S2). Post Hoc tests results are indicated on the graph (**, p ≤ 0.01; ***, p ≤ 0.001).

In order to verify if the difference of the body weight observed in hypothyroid mice was due to a lower food consumption induced by PTU treatment, we measured the food intake of each group (Fig.2B-S1). Interestingly, we first observed a difference in food intake between the two euthyroid groups according to the strain (p < 0.0001). Euthyroid WSB/EiJ mice have a higher food intake than C57BL/6J mice despite their lean phenotype, reflecting the efficient metabolic character of this strain. Besides, both euthyroid C57BL/6J and WSB/EiJ mice maintained their food intake throughout the weeks of treatment (p > 0.05), although the food intake of euthyroid C57BL/6J mice was reduced the very last week of the treatment (probably because mice reached a stable body weight, p ≤ 0.05). However, PTU treatment significantly reduced the food intake in both hypothyroid C57BL/6J and WSB/EiJ mice, compared to their euthyroid control groups (p < 0.05). For both hypothyroid groups, the food intake was reduced from the first week of treatment (p ≤ 0.05) and was overall maintained throughout the treatment (p > 0.05). Therefore, the significant weight-loss observed in hypothyroid C57BL/6J mice and not in hypothyroid WSB/EiJ mice, was not due to a different food intake as both strains maintained their food consumption during the treatment. Furthermore, the different weight was either not due to growth impairment as assessed by femur length measurements which revealed no differences between groups (Fig.S2). Although euthyroid WSB/EiJ mice consumed more food than C57BL/6J mice related to their weight, their eWAT weight was considerably (2-times) lower than C57BL/6J mice (p = 0.003; Fig.2C), highlighting the low-fat mass in WSB/EiJ mice at the basal state. In response to PTU-treatment, eWAT weight was unchanged in C57BL/6J mice (p >0.05) whereas decreased in WSB/EiJ mice (p = 0.007). Since leptin is known to be proportional to body fat mass (Fig. 2D), circulating leptin levels were accordingly lower in euthyroid WSB/EiJ mice compared to C57BL/6J mice (p = 0.0001), reflecting here again the lean phenotype of WSB/EiJ strain. In response to PTU treatment, circulating leptin levels were decreased in C57BL/6J mice (p = 0.0003) whereas levels were maintained in WSB/EiJ mice (p >0.05), reflecting the weight loss of C57BL/6J mice but not for WSB/EiJ mice. However, leptin mRNA expression in the eWAT (Fig.2E), where leptin is principally produced, was not altered in both hypothyroid mouse strains (p >0.05). These results showed that PTU-treatment altered differentially the body weight and the fat mass between both strains.

#### 2.2. Circulating lipids

Circulating lipid profiles were assessed and revealed a differential effect of hypothyroidism between both mouse strains (Fig.3), consistent with fat mass results. Triglycerides (TGs) and non-esterified fatty acid (NEFA) levels were decreased in hypothyroid C57BL/6J mice (p < 0.01), whereas levels remained unchanged in hypothyroid WSB/EiJ mice (p > 0.05; Fig.3A-B). These results were in line with the difference of the body weight observed between the two hypothyroid strains. However, both C57BL/6J and WSB/EiJ mice strain exhibited an increase of cholesterol levels in response to PTU (p < 0.001; Fig.3C), which were even more pronounced in WSB/EiJ mice (by 29% compared to hypothyroid C5BL/6J mice; p < 0.001). Low-density lipoprotein (LDL) levels, which carried cholesterol into the blood, followed the same profile as cholesterol levels for both hypothyroid strains (p < 0.001: Fig.3D). Similarly, high-density lipoprotein (HDL) levels, which export cholesterol from blood to the liver for clearance, increased in both hypothyroid mouse strains and more importantly in WSB/EiJ mice (p < 0.001; Fig.3E). Therefore, both mouse strains exhibited hypercholesterolemia in response to hypothyroidism. In contrast, TGs and NEFA levels were reduced in hypothyroid C57BL/6J mice whereas maintained in WSB/EiJ mice, indicating an alteration of lipid content only in hypothyroid C57BL/6J mice.

**Figure 3.**
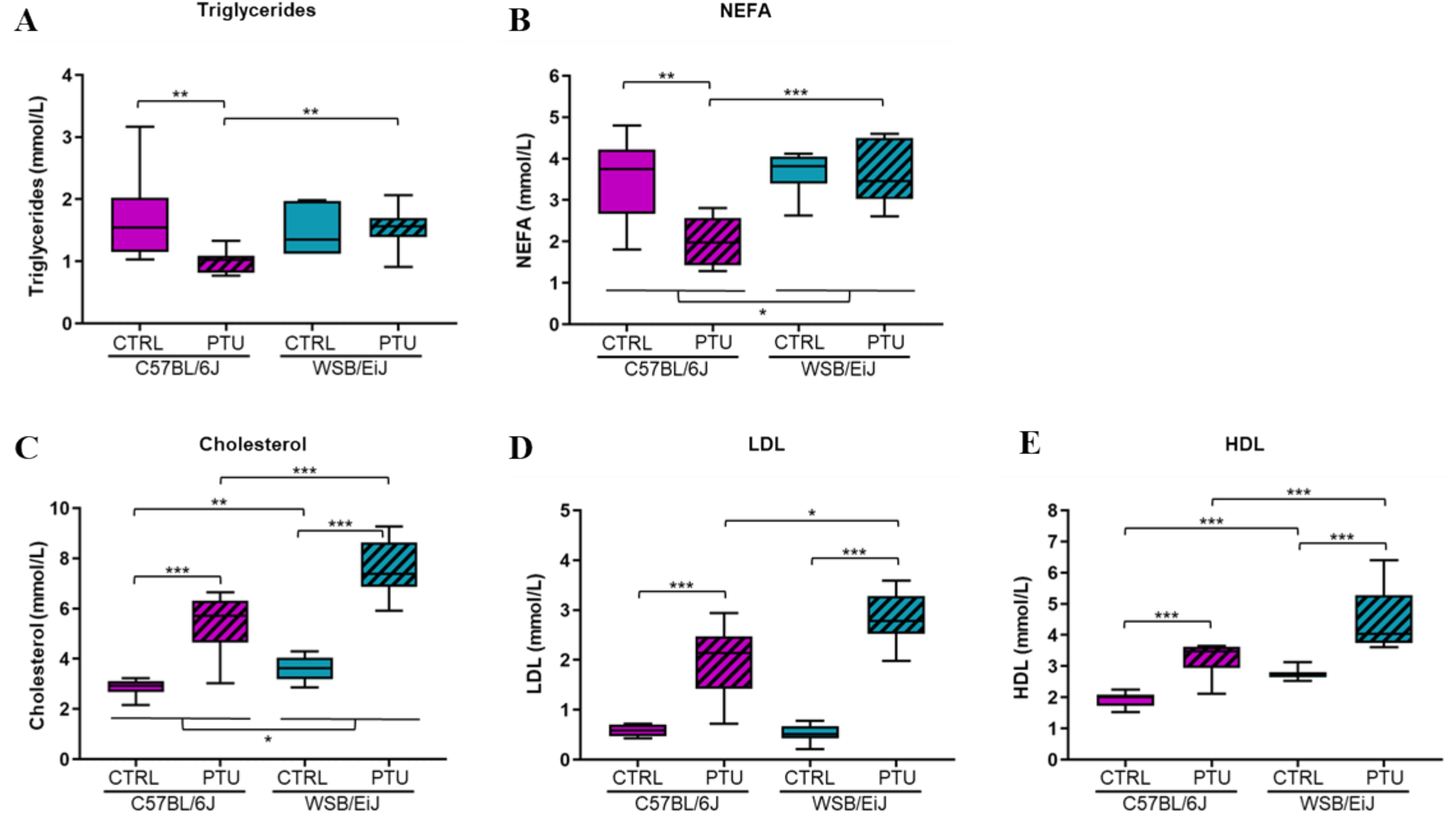
Circulating lipid patterns are altered differentially between mouse strains in response to hypothyroidism. **(A)** Triglycerides (in mmol/L) and **(B)** non-esterified fatty acid (NEFA in mmol/L) levels were decreased in hypothyroid C57BL/6J mice whereas unchanged in WSB/EiJ mice. **(C)** Cholesterol (in mmol/L), **(D)** low density lipoprotein (LDL in mmol/L) and **(E)** high density lipoprotein (HDL in mmol/L) levels were increased in both mouse strains after PTU-treatment. Boxplot represents median values and min-max whiskers. Post Hoc tests results are indicated on the graph (n = 7-8 per group; non-parametric two- and one-way ANOVA with permutations test; *, p ≤ 0.05; **, p ≤ 0.01; ***, p ≤ 0.001).

#### 2.3. Hepatic and adipose lipid metabolism

Next, to better understand the differential lipid responses to hypothyroidism between the two strains, we investigated whether peripheral lipid metabolism was altered by PTU treatment. Thus, we quantified by RT-qPCR the expression of lipid-related genes in the liver, the main tissue in which lipid synthesis occurs (Fig.4). Peripheral hypothyroidism was first confirmed by the expression of *Dio1*, a key marker of peripheral hypothyroidism (Bianco and Kim, 2006), which was drastically downregulated in both hypothyroid strains compared to their euthyroid group (p < 0.05; Fig.4A). The expression of lipogenic genes (*Fasn, Acacα*), key actors of the lipogenesis pathway by which TGs are synthetized, were downregulated by 2-fold in both hypothyroid mouse strains compared to the untreated mice (Fig.4B-C) (p < 0.01, see table S2). Furthermore, the transcription factor *Chrebp* which is involved in the regulation of this pathway, was also downregulated in both hypothyroid strains compared to their euthyroid group (“Treatment” effect p < 0.001; Fig.4D). The expression of *Pparα*, implicated in hepatic lipid metabolism and utilization, was downregulated in both hypothyroid strains compared to their euthyroid control (“Treatment” effect p < 0.001; Fig.4E). However, expression of *Ppargc1α*, known as a co-activator of *Pparα* and essential for fatty acid oxidation, was upregulated in C57BL/6J mice in response to PTU treatment (p < 0.01) but remained unaffected in WSB/EiJ mice (p > 0.05; Fig.4F). Another key actor of fatty acid oxidation, *Fgf21*, was also upregulated in hypothyroid C57BL/6J mice but slightly downregulated in WSB/EiJ mice (0.01 < p < 0.05; Fig.4G). Taken together, these results showed that PTU-treatment impaired the hepatic lipid metabolism in both strains by globally reducing TGs synthesis.

**Figure 4.**
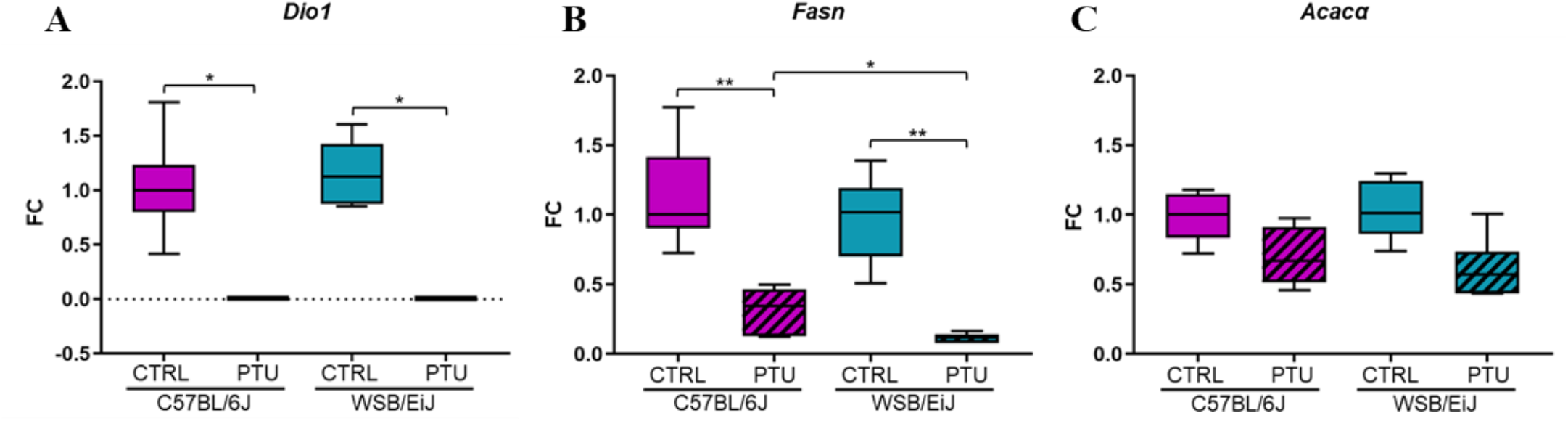
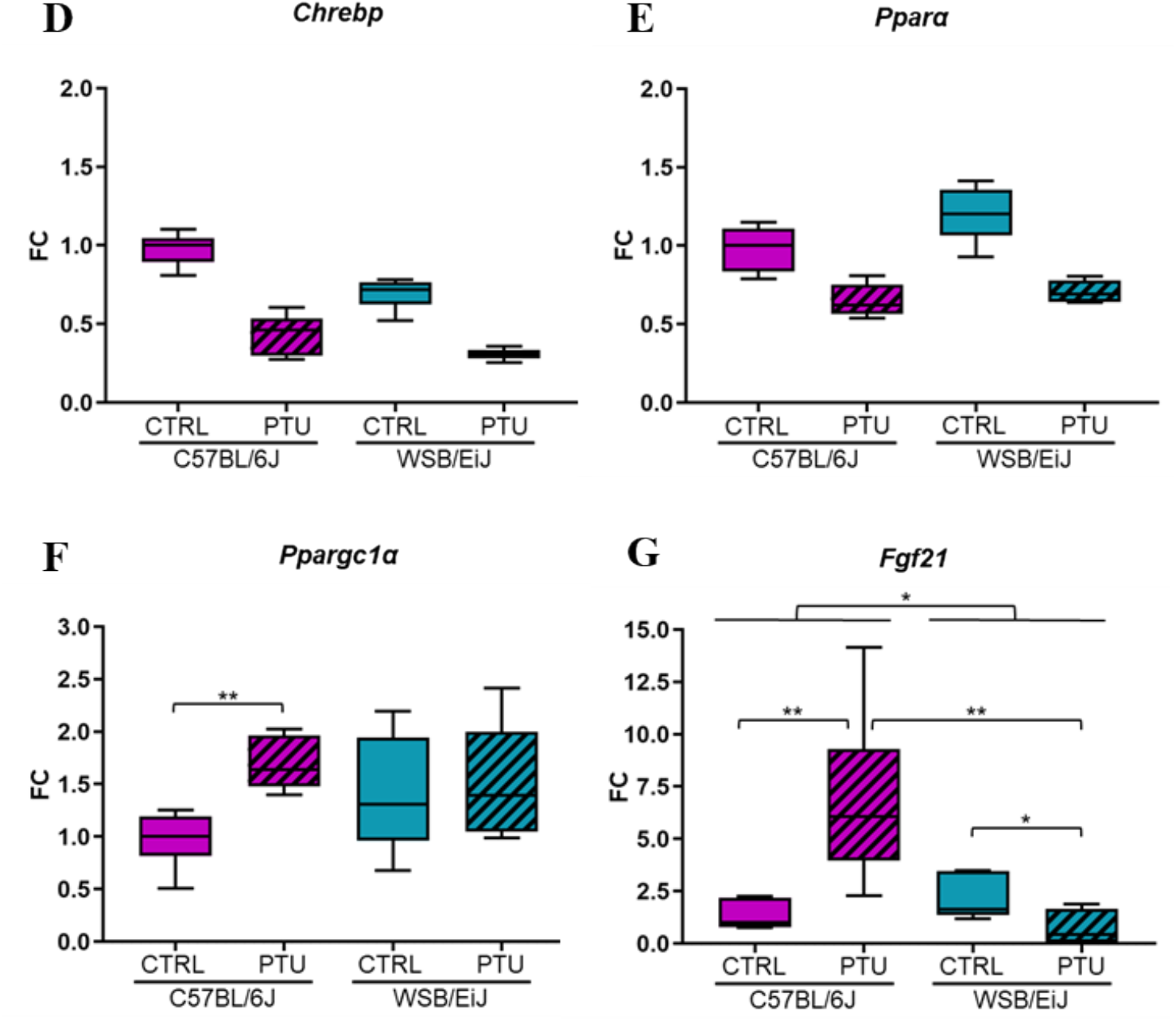
Hepatic lipid metabolism is reduced in both strains in response to hypothyroidism. **(A)** Liver expression of *Dio1* gene was totally down-regulated in C57BL/6J and WSB/EiJ mice after 7 weeks of PTU-treatment. Expression of lipogenesis key genes *Fasn* **(B)** and *Acacα* **(C)** involved in the fatty acid and TGs synthesis (referred as de novo lipogenesis (DNL)) were downregulated in both strains after PTU-treatment, as well as *Chrebp* **(D)**, the DNL-regulated transcription factor. Expression of *Pparα* **(E)**, *Ppargc1α* (**F)** and *Fgf21* (**G)** genes, involved in lipid metabolism regulation and fatty acid oxidation, were differentially regulated between strains. Data are represented as relative fold-change expression (FC). Boxplot represents median values and min-max whiskers. Post Hoc tests results are indicated on the graph (n = 5-6 per group; non-parametric two-way ANOVA with permutations test; *, p ≤ 0.05; **, p ≤ 0.01).

Next, we were intrigued to know by which mechanisms hypothyroid WSB/EiJ mice maintained their circulating lipid contents, as TGs synthesis was reduced in the liver. For this purpose, we quantified the expression of lipid-related genes in the eWAT which is another important source of lipids (Fig.5). Surprisingly, we observed that the expression of *Fasn* and *Acacα* remained unchanged in hypothyroid C57BL/6J mice (p < 0.05) whereas upregulated by almost 3-fold in hypothyroid WSB/EiJ mice (p < 0.05; Fig.5A-B), suggesting an enhanced lipogenesis only in WSB/EiJ mice in response to PTU treatment. Although lipogenesis is regulated by the transcription factor *Pparγ*, its expression was unchanged in both hypothyroid strains (p > 0.05; Fig.5C). Another potential source of lipids is the lipolysis which catalyzes TGs (stored in the form of lipid droplets) into free fatty acid (FFA) by PNPLA2 enzyme in the eWAT. In response to PTU-treatment, the mRNA expression of this key-enzyme was unaffected in C57BL/6J mice (p > 0.05) whereas upregulated in hypothyroid WSB/EiJ mice (by 110% compared to the euthyroid group, p > 0.05; Fig.5D). However, the difference observed in hypothyroid WSB/EiJ mice did not reach the significance due to the high variability of this group. Besides, the expression of *Ppargc1a*, the master-regulator of fatty acid oxidation and thermogenesis in the eWAT, was quantified as body weight and lipid metabolism was dysregulated in hypothyroid mice of each strain (Fig.5E). We observed that *Ppargc1a* mRNA expression was downregulated in both hypothyroid groups (p < 0.01), and more importantly (2-fold compared to euthyroid) in hypothyroid C57BL/6J mice compared to WSB/EiJ mice (p < 0.001). Consequently, lipid metabolism in eWAT was improved in WSB/EiJ mice in response to PTU treatment and not in C57BL/6J mice, which could explain sustained circulating lipid levels in the WSB/EiJ strain. However, C57BL/6J mice were more affected by hypothyroidism than WSB/EiJ mice which developed mechanisms to compensate for lipid deficiency.

**Figure 5.**
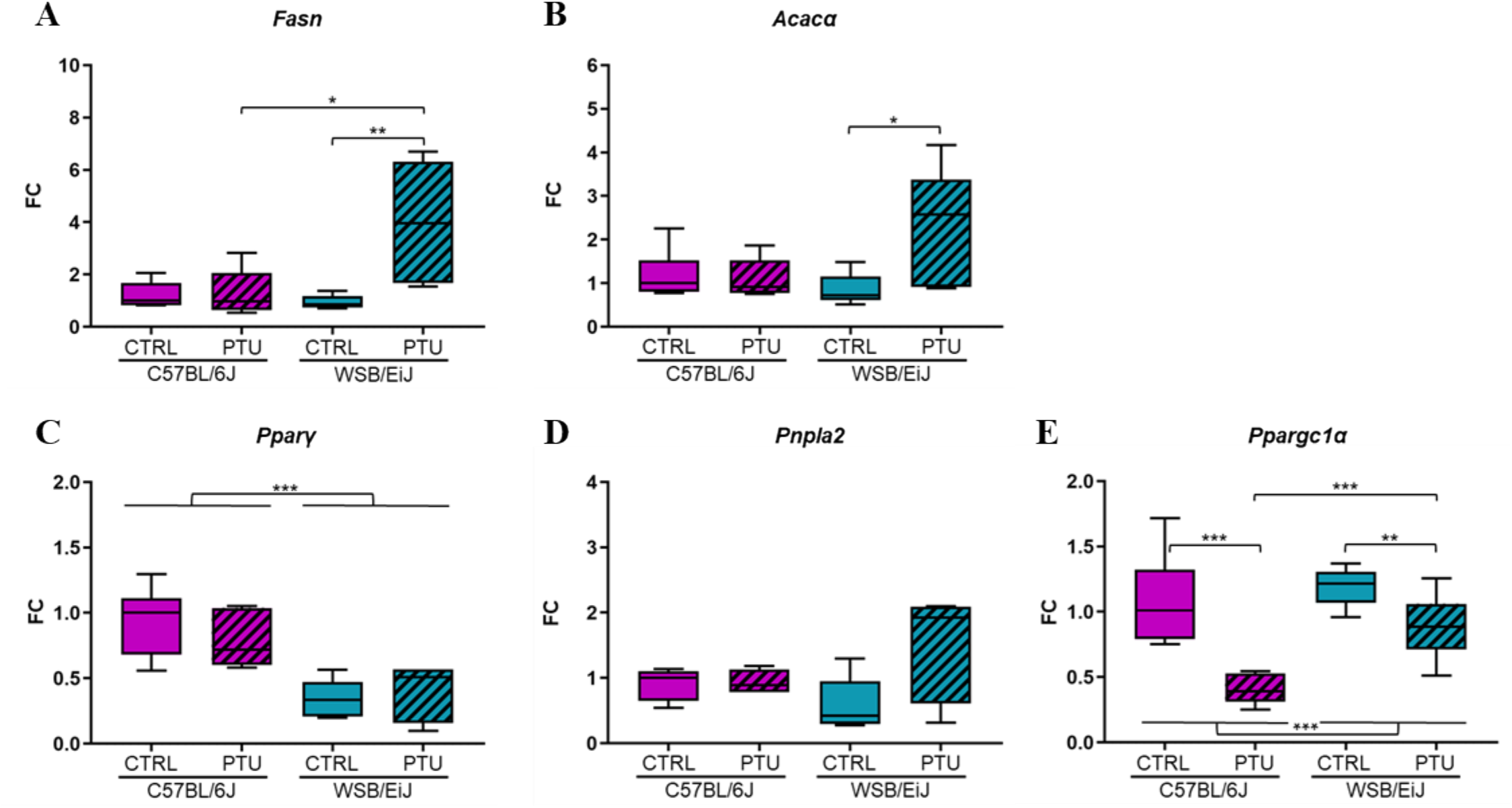
Adipose lipid metabolism is globally enhanced in WSB/EiJ mice and unchanged in C57BL/6J mice under PTU-treatment. Expression in the eWAT of lipogenesis key genes *Fasn* **(A)** and *Acacα* **(B)** involved in the fatty acid and TGs synthesis were upregulated only in WSB/EiJ mice after PTU-treatment. **(C)** Expression of the DNL-regulated transcription factor *Pparγ* gene was different between strains. **(D)** Expression of *Pnpla2* transcript, catalyzing TGs into fatty acids (referred as lipolysis) seemed to be increased only in WSB/EiJ mice but did not reach significance. Expression of *Ppargc1α* gene, involved in fatty acid oxidation, was downregulated between strains. Data are represented as relative fold-change expression (FC). Boxplot represents median values and min-max whiskers. Post Hoc tests results are indicated on the graph (n = 5-6 per group; non-parametric two-way ANOVA with permutations test; *, p ≤ 0.05; **, p ≤ 0.01).

#### 2.4. Hypothalamic regulation of energy balance

We further investigated the effect of hypothyroidism on the hypothalamic expression of genes involved in the control of food intake and body weight (Fig.6). We evaluated particularly the expression of orexigenic (*Agrp*) and anorexigenic factors (*Pomc, Mc4R, LepR*). We observed that the expression of *Agrp* was upregulated by 138% and 83% in hypothyroid C57BL/6J (p < 0.05) and WSB/EiJ mice, respectively, compared to their euthyroid group, but the difference did not reach the significance for WSB/EiJ mice due to the high variability of this group (p > 0.05; Fig.6A). In contrast, the expression of anorexigenic factors were downregulated in hypothyroid C57BL/6J mice after PTU treatment compared to controls (p < 0.05; Fig.6B-D, see table S2). However, the expression of *Mc4r* and *Lepr* were maintained (p > 0.05) whereas *Pomc* expression was decreased in hypothyroid WSB/EiJ mice compared to their control group (p < 0.05). Taken together, these results showed that hypothyroidism stimulates the hypothalamic orexigenic circuit in both strains. However, unchanged *Mc4r* and *LepR* expression in hypothyroid WSB/EiJ mice strain reflected their adaptability to maintain their body weight, contrary to C57BL/6J mice.

**Figure 6.**
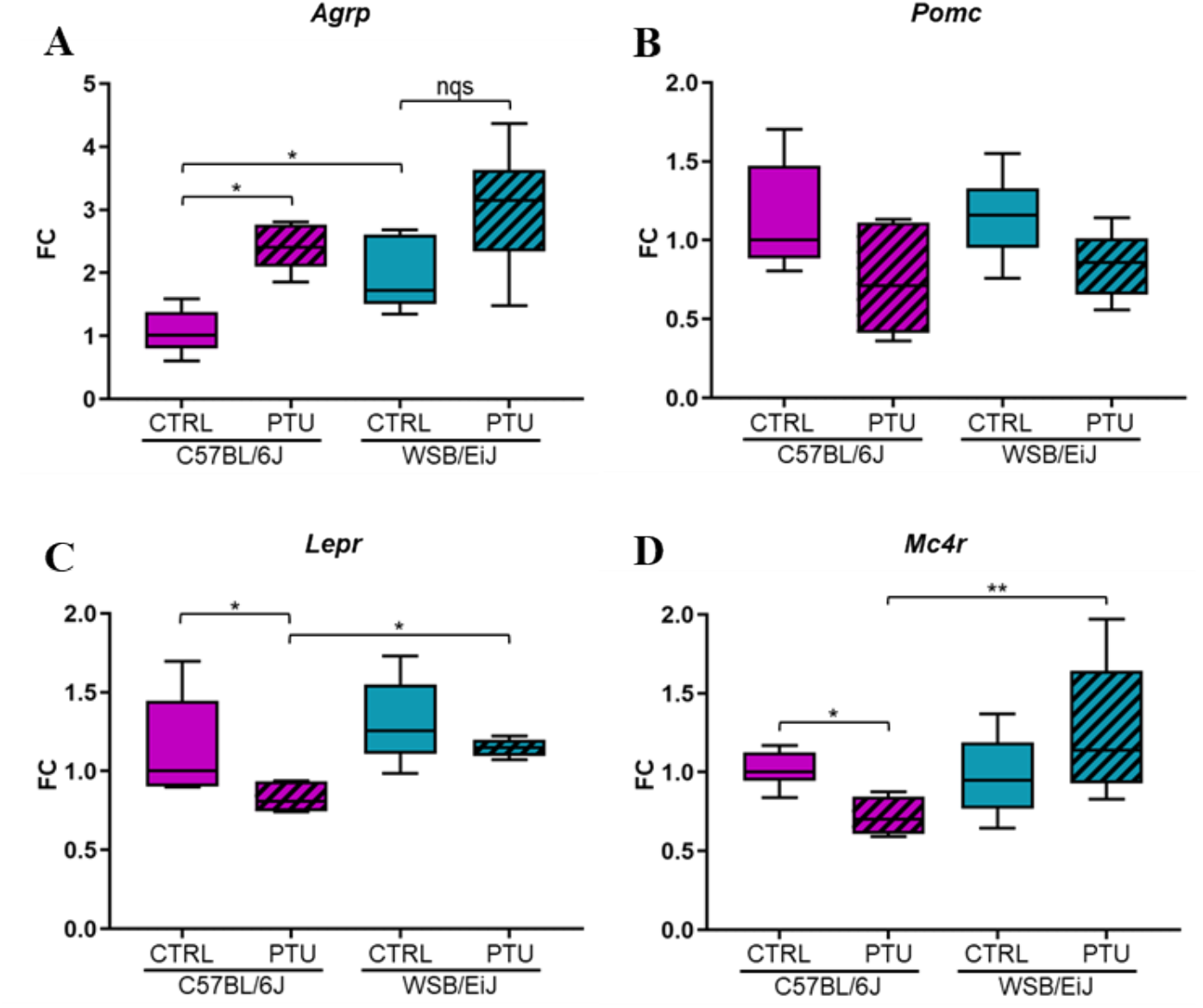
PTU-treatment effects on hypothalamic energy balance. Expression of appetite-stimulating (orexigenic) neuropeptide *Agrp* **(A)**, appetite-suppressing (anorexigenic) neuropeptide *Pomc* **(B)**, melanocortin 4 receptor *Mc4r* **(C)** and leptin receptor *Lepr* **(D)** in response to 7-weeks of treatment. Data are represented as relative fold-change expression (FC). Boxplot represents median values and min-max whiskers. Post-Hoc tests results are indicated on the graph (n = 5-6 per group; non-parametric one- and two-way ANOVA with permutations test; *, p ≤ 0.05; **, p ≤ 0.01; ***, p ≤ 0.001; nqs: not quite significant).

### III. Hypothyroidism consequences on peripheral and hypothalamic inflammation

#### 3.1. Peripheral inflammation

Because lipid metabolism was dysregulated mainly in hypothyroid C57BL/6J mice, we questioned whether an inflammatory response was developed within the two mouse strains (Fig.7). The expression of pro-inflammatory (*Tnfα, Il6, Il1β*) and anti-inflammatory (*Il10*) genes were first evaluated in the eWAT, an active endocrine organ producing and secreting a plethora of cytokines and hormones ensuring homeostasis (Coelho, Oliveira and Fernandes, 2013) (Fig.7A). The expression of these inflammatory genes was overall downregulated in hypothyroid C57BL/6J mice (p < 0.05) whereas unchanged in WSB/EiJ mice (p > 0.05) compared to the untreated mice. These results were confirmed by circulating cytokine measurements that showed similar profile patterns (Fig.7B). The cytokine levels were overall lower in WSB/EiJ mice than in C57BL/6J mice according to the strain (p < 0.05). In response to PTU-treatment, serum IL1β and IL10 levels were decreased in C57BL/6J mice (by 77% and 27% respectively compared to their euthyroid group; p < 0.05). Inversely, IL1β levels were increased in WSB/EiJ mice (by 110% compared to their control group, p < 0.05) whereas IL10 levels were unaffected in response to hypothyroidism (p < 0.05). IFNγ and IL-6 levels were remained unchanged in both mouse strains in response to PTU treatment (p > 0.05). Taken together, PTU treatment induced differential peripheral inflammatory responses between strains, that were reduced in C57BL/6J mice and maintained in WSB/EiJ mice.

**Figures 7.**
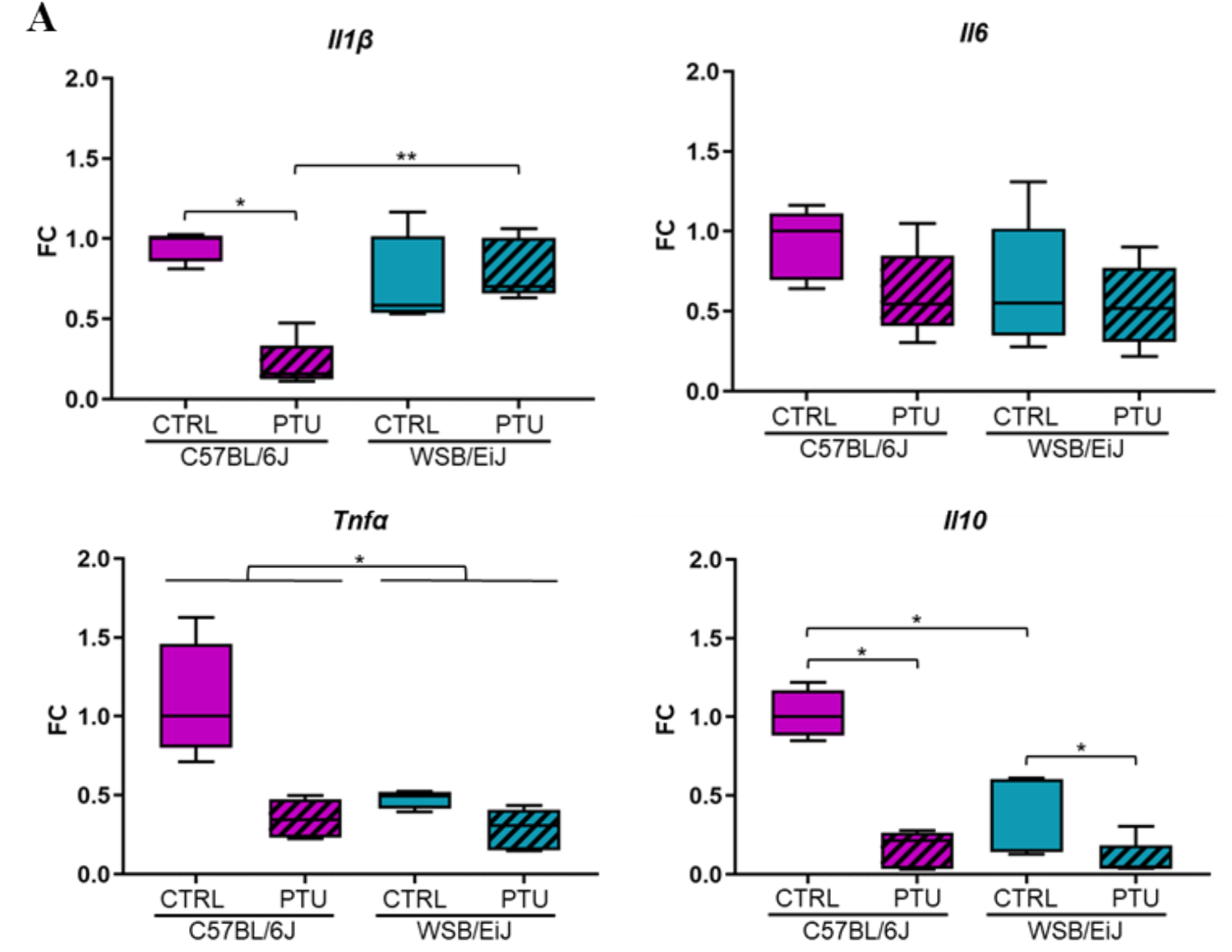
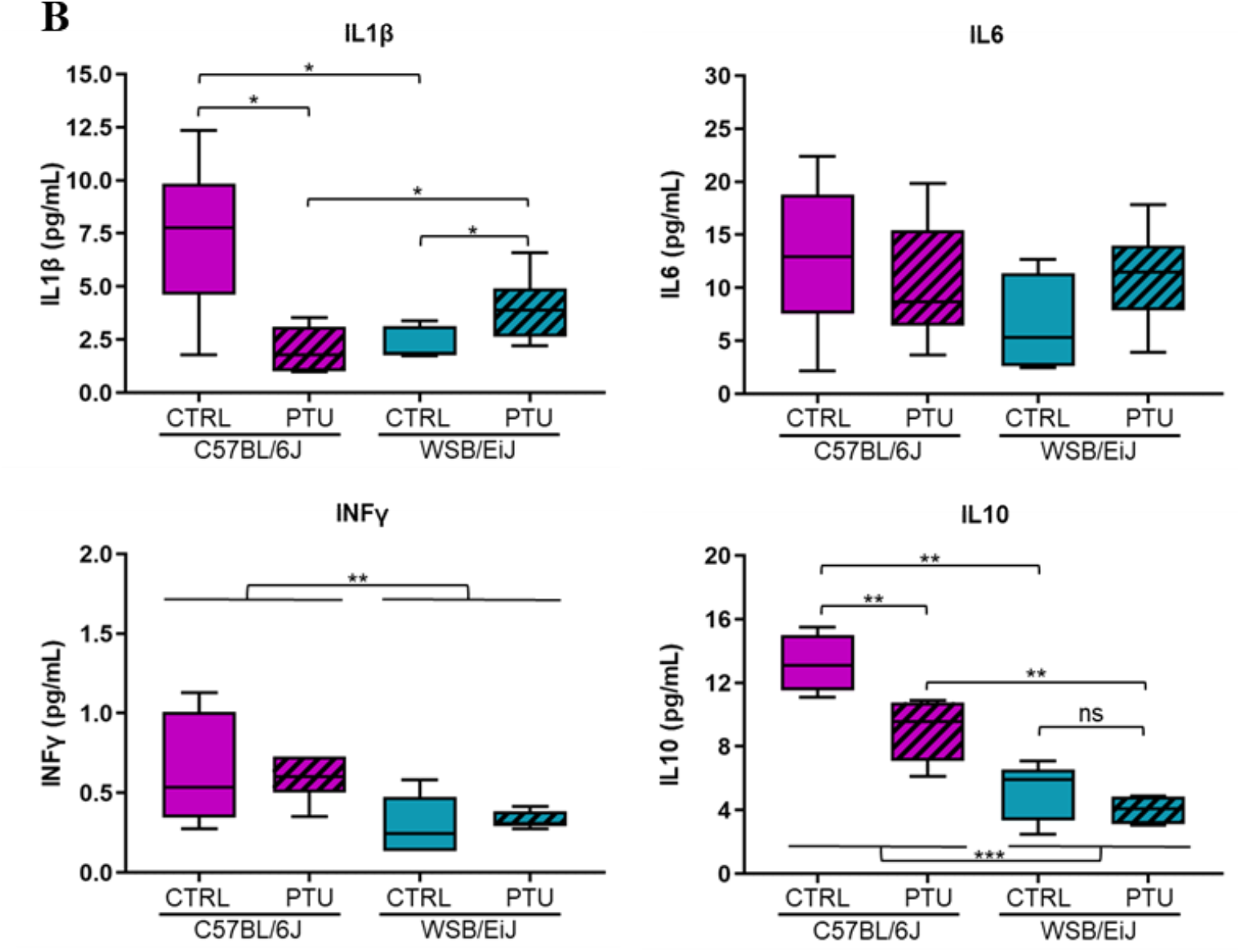
Effect of PTU-treatment on the inflammatory cytokines. **(A)** Expression of pro-inflammatory *Il1B, Tnfα, Il6* and anti-inflammatory *Il10* cytokine genes in the eWAT were overall downregulated in hypothyroid C57BL/6J mice whereas unchanged in WSB/EiJ mice. Data are represented as relative fold-change expression (FC). **(B)** Circulating inflammatory cytokines measured by ELISA: IL10 (in pg/mL), IL1β (in pg/mL), IFNγ (in pg/mL), IL6 (in pg/mL) levels were overall decreased in hypothyroid C57BL/6J mice whereas unchanged in WSB/EiJ mice. Boxplot represents median values and min-max whiskers. Post Hoc tests results are indicated on the graph (n = 5-6 per group; non-parametric two- and one-way ANOVA with permutations test; *, p ≤ 0.05; **, p ≤ 0.01; ***, p ≤ 0.001).

#### 3.2. Hypothalamic inflammation

We also investigated central inflammation by quantifying astrocytes and microglia populations as inflammatory markers. Hypothalamic glial cells were first checked in the ARC (Fig.8). In agreement with peripheral inflammatory markers, density of IBA1+ microglia was slightly decreased in hypothyroid C57BL/6J mice (by 14% compared to their euthyroid control, p < 0.05), whereas maintained in hypothyroid WSB/EiJ mice (p > 0.05) (Fig.8A-B). However, density of GFAP+ astrocytes was unaffected by PTU treatment in both strains (“Treatment” effect p > 0.05) (Fig8.C-D) but was overall lower in WSB/EiJ mice than in C57BL/6J mice indicating low inflammatory markers in this strain. Therefore, these results showed that inflammatory markers were reduced in C57BL/6J mice whereas maintained in WSB/EiJ mice in response to hypothyroidism, consistent with the reduction of peripheral lipid metabolism in C57BL/6J strain.

**Figure 8.**
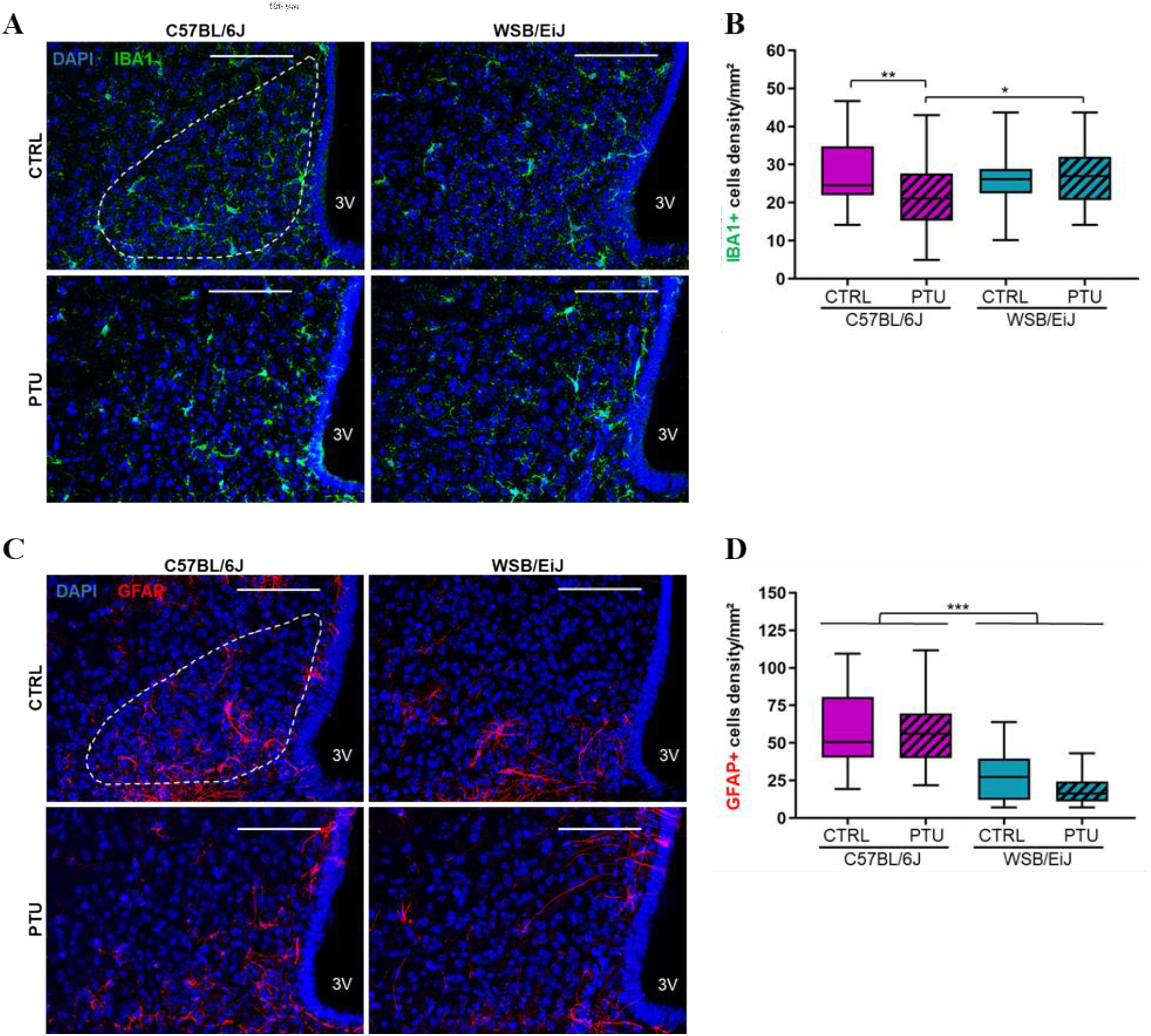
Effect of hypothyroidism on glial cells density in the hypothalamic arcuate nucleus (ARC) of mouse strains. **(A)** Representative confocal images of IBA1+ microglia (green), in the ARC (white ROI) of euthyroid (CTRL) and hypothyroid (PTU) C57BL/6J (left panel) and WSB/EiJ (right panel) mice. Cells nuclei are stained with DAPI (in blue) **(B)** Quantitative analysis revealed a decrease of microglia density in the ARC of C57BL/6J mice whereas unchanged in WSB/EiJ mice in response to PTU-treatment **(C)** Representative confocal images of GFAP+ astrocytes (red), in the ARC of euthyroid (CTRL) and hypothyroid (PTU) C57BL/6J (left panel) and WSB/EiJ (right panel) mice. Cells nuclei are stained with DAPI (in blue) **(D)** Density of GFAP+ astrocytes remained unchanged in the ARC of both mouse strains after PTU-treatment and a strain effect revealed less astrocytes density in the ARC of WSB/EiJ compared to C57BL/6J mice. Boxplot represents median values and min-max whiskers. Strain effect result is indicated on the graph (n = 4 mice per group, n = 2-6 sections per mouse and n= 4 ROI per section; non-parametric two-way ANOVA with permutations test; *, p ≤ 0.05; **, p ≤ 0.01; ***, p ≤ 0.001). Scale bars = 100μm.

## Discussion

Hypothyroidism being induced by treating both mouse strains with PTU, we therefore assessed the thyroid status of mice. 7-weeks of PTU treatment induced hypothyroidism in both strains, as confirmed by the drastic reduction of serum TH levels in both mouse strains.

HPT axis is key actor in the regulation of energy balance and hypothyroidism is well described to cause metabolic dysregulations in rodents and humans (Song, Yao and Ying, 2011; Mullur, Liu and Brent, 2014). In response to hypothyroidism, the metabolic dysregulations were first manifested by weight loss in both mouse strains comparing to their euthyroid control. This was similarly observed in hypothyroid rodents treated with PTU or MMI starting from 2 weeks of treatment (Decherf *et al*., 2010; Chaalal *et al*., 2014; Herwig *et al*., 2014). This weight-loss induced by hypothyroidism is probably due, in part, to an alteration of food intake mechanisms as both strains reduced their consumption when treated with PTU comparing to euthyroid mice, starting from the 1^st^ week. This was also observed in hypothyroid rodents associated to a weight loss (Decherf *et al*., 2010; Herwig *et al*., 2014). Conversely, an increase in food intake and weight gain was observed in hyperthyroid mice treated with T4 for 9 weeks (Rakov *et al*., 2017). Therefore, deregulation of thyroid axis could have effect on appetite and body weight, as THs are key regulators of energetic metabolism. However, in response to PTU treatment, a differential body weight variation was observed between hypothyroid mice of both strains independently of the food intake as it was maintained throughout the treatment: hypothyroid WSB/EiJ mice maintained their body weight throughout the treatment, whereas hypothyroid C57BL/6J mice showed a weight-loss of 5% that was pronounced at week five of the treatment. The differential body weight between the two strains was reflected by leptin concentrations, known to be secreted by the white adipose tissue (WAT) in proportion to body fat (Maffei *et al*., 1995). The reduced serum leptin levels in hypothyroid C57BL/6J mice reflected their reduced body fat, whereas these levels were maintained in WSB/EiJ mice accordingly to their stable body weight in response to treatment. Moreover, in hypothyroid C57BL/6J mice, free fatty acids (FFAs) and triglycerides (TGs) levels were decreased whereas maintained in hypothyroid WSB/EiJ mice, revealing differential hypothyroidism effect on the lipid machinery between both strains. These lipid concentrations are concordant with the weight differences as well as the leptin levels observed between hypothyroid mouse strains. Numerous studies have also reported a decrease in lipid levels in response to hypothyroidism. In mice treated with PTU for 7 weeks, a decrease in serum TGs and FFA levels was observed (Olichwier *et al*., 2021) ; a decrease in plasma TGs concentrations associated with a reduction in fat mass, were reported in rats treated with MMI for 22 days (Herwig *et al*., 2014). Taking together, these data confirm that PTU-treatment induced alteration in lipid metabolism, particularly in C57BL/6J mice. In contrast, WSB/EiJ mice showed a phenotype of resistance to lipid alteration induced by hypothyroidism. However, hypothyroidism induced an impairment of cholesterol metabolism as both strains exhibited an increase of cholesterol circulating levels together with an increase of HDL and LDL lipoproteins (Abbas *et al*., 2008; Rizos, 2011; Sinha, Singh and Yen, 2018). This raises the question of how hypothyroidism altered the hepatic cholesterol metabolism in both strains since THs control cholesterol anabolic and catabolic processes including the reverse cholesterol transport mechanism in the liver (Ritter, Amano and Hollenberg, 2020). THs are critical actors of the peripheral metabolism by exerting direct effects on metabolically active organs such as liver and WAT (Mullur, Liu and Brent, 2014). It has been extensively described in humans and rodents that hypothyroidism decreased the carbohydrate and lipid metabolism leading to an insulin-resistance state (Rizos, 2011; Mcaninch and Bianco, 2014; Duntas and Brenta, 2018). FFAs and TGs are the main lipid components produced particularly in the liver. THs are a well-known activator of de novo hepatic lipogenesis (DNL) by which the FFAs are synthetized, as they directly stimulate the transcription of several key genes such as *Fasn* and *Acaca*. THs control as well the expression of transcription factors involved in the DNL such as *Chrebp* (Sinha, Singh and Yen, 2018; Ritter, Amano and Hollenberg, 2020). After their synthesis, FFAs are esterified, accumulated as TGs and released in blood circulation. In this present study, hypothyroidism induced an impairment of the lipid metabolism which was first reflected by the decrease of hepatic lipid synthesis. This was revealed by the downregulation of the lipogenic genes expression, mainly *Fasn, Acacα* and *Cherbp*, in the liver of both strains, suggesting a reduced hepatic DNL and thus lipid synthesis. Similar results were previously observed in hypothyroid rodents (Ferrandino *et al*., 2017; Gnoni *et al*., 2019). However, eWAT DNL was differentially altered between both hypothyroid strains in response to the hepatic lipid deficiency. Adipocyte DNL is an important source of endogenous lipids and plays crucial roles in maintaining systemic metabolic homeostasis. The biosynthesis of TGs in the eWAT results from FFA synthesis by DNL in case of fat excess. Inversely, lipolysis is the catabolic process leading to the breakdown of TGs by adipose triglyceride lipases as PNPLA2 to FFAs that can be released and delivered to the liver when energy fuel is required (Shi and Burn, 2004; Luo and Liu, 2016). Both adipocyte lipogenesis and lipolysis are regulated by THs (Obregon, 2014; Song, Xiaoli and Yang, 2018). In C57BL/6J mice, hypothyroidism had no effect on either adipose lipogenesis nor lipolysis as the expression of respective genes (*Fasn, Acacα, Pnpla2*) remained unchanged. In contrast, adipose lipogenic and lipolytic genes were considerably upregulated in hypothyroid WSB/EiJ mice, suggesting an enhanced lipid mobilization in eWAT in response to the reduced hepatic lipid machinery. Our results demonstrated that hypothyroidism reduced hepatic lipid metabolism in both strains but was rescued only in WSB/EiJ mice by increasing lipid availability from adipocytes. This could explain the reduced circulating lipid levels in hypothyroid C57BL/6J mice whereas maintained in WSB/EiJ mice. Furthermore, eWAT weight was decreased only in hypothyroid WSB/EiJ mice which could result from hydrolyzed fat occurred in this tissue, highlighting here again the metabolic flexibility of WSB/EiJ mice. Taken together these results indicate that hypothyroidism reduced hepatic lipid biosynthesis in both strains, which was compensated only in WSB/EiJ mice by stimulating lipid mobilization from eWAT.

In metabolic dysregulations such as obesity, a lowgrade inflammatory response is triggered by eWAT which releases cytokines and chemokines (Masoodi *et al*., 2015; Guillemot-legris and Muccioli, 2017). In this study, we were interested to know if metabolic dysregulations induced by hypothyroidism promote peripheral inflammation. The inflammatory response was overall reduced in C57BL/6J mice whereas maintained in WSB/EiJ mice in response to PTU-treatment. Indeed, pro-inflammatory and anti-inflammatory cytokine transcripts, such as *Tnfa, Il1b* and *Il10* respectively, were downregulated in eWAT hypothyroid C57BL/6J whereas maintained in WSB/EiJ mice, in parallel with circulated cytokine levels. Inversely to the consequences effect of lipid overload (Guillemot-legris and Muccioli, 2017), we speculated that lipid deficiency and the weight loss occurring in hypothyroid C57BL/6J mice could explain, in part, the repression of the inflammatory responses in this strain. Indeed, the absence of inflammatory response could be explained by the reduction of lipid level known to stimulate pro-inflammatory cytokines (Ertunc and Hotamisligil, 2016). Corroborating other findings, reduction in body fat is positively correlated with the immune function in many species including humans (Norgan, 1997; Carlton, Demas and French, 2012). However, maintenance of inflammatory reaction in WSB/EiJ mice could result from their compensatory mechanisms deployed to overcome THs deficiency. This highlight here again the efficiency of this strain to sustain homeostasis despite peripheral hypothyroidism.

In the brain, THs are key regulators of glial cells, particularly astrocytes and microglia which are crucial for neuronal protection and brain homeostasis (Fields and Stevens-Graham, 2002). Moreover, glial cells are considered as brain immune cells, particularly the microglia, since they promote inflammatory processes in inflammatory-related diseases such as obesity and neurodegenerative diseases (Guillemot-legris and Muccioli, 2017; Kempuraj *et al*., 2017; Walker *et al*., 2020). In the hypothalamus, these two glial cell populations are highly sensitive to metabolic deregulations and act as peripheral metabolic sensors mainly in the mediobasal hypothalamus (including the hypothalamic arcuate nucleus (ARC)) considered as an interface between the CNS and the periphery (García-Cáceres *et al*., 2012; Kälin *et al*., 2015; Terrien *et al*., 2019). In our study, hypothyroidism altered microglia density as it was reduced only in the ARC of C57BL/6J mice (in the range of 14%) whereas astrocytes density was unaffected in both strains. The decrease in microglia population was also reported in hypothyroid neonatal rat since THs control microglia development and maturation (Mohácsik *et al*., 2011). We presumed that the decrease in hypothalamic microglia density in hypothyroid C57BL/6J mice could result from the reduction of peripheral inflammation in response to declined lipid metabolism induced by hypothyroidism.

Besides, the hypothalamus is the key regulator of energy balance and body weight maintenance (Schwartz, 2006; Roh and Kim, 2016). It involves the coordination of two main hypothalamic neuronal circuits: orexigenic and anorexigenic circuits particularly AGRP / NPY and POMC / MC4R neurons, respectively (Flier, 2004; Schwartz, 2006). Moreover, metabolic signals, as leptin, orchestrate the regulation of these neuronal circuits in order to modulate the food intake. In response to hypothyroidism, mRNA levels showed a stimulation of the orexigenic neurons (*Agrp*) and an inhibition of the anorexigenic neurons (*Pomc*), suggesting a coordinated response observed in both strains in order to increase food intake to restore the stocks. Indeed, the stimulation of the orexigenic neurons could be associated with the decrease of leptin levels in hypothyroid C57BL/6J as lipid synthesis was reduced. Therefore, we observed the classical leptin action as a negative feedback loop in hypothyroid C57BL/6J mice to promote energy balance and the recovery of the weight loss. The combination of these responses has been well demonstrated in diet induced weight loss conditions such as starvation and caloric restriction (Ahima, 2008; Jensen *et al*., 2013). However, leptin levels were maintained in hypothyroid WSB/EiJ mice as body weight was maintained. This presumes that stimulation of orexigenic circuits in WSB/EiJ mice is not dependent on leptin signaling, as observed in C57BL/6J mice, and underlies the involvement of other homeostatic mechanisms such as melanocortin pathway. Anyhow, we supposed that the activation of orexigenic pathways in response to hypothyroidism could explain the maintenance of food consumption in both hypothyroid mice strain throughout the treatment. Despite, their consumption remains lower than that of euthyroid mice, suggesting clear effect of hypothyroidism on the control of appetite. Hypothalamic melanocortinergic pathway is known to be involved in the regulation of appetite and weight maintenance (Koch and Horvath, 2014). It is triggered when POMC is activated and AGRP/NPY is inhibited, thus mediating anorexigenic signals through melanocortin receptors (MC4R/MC3R) (O’Rahilly,

Yeo and Farooqi, 2004; Vella *et al*., 2011). As orexigenic circuit was stimulated in hypothyroid mice, the expression of *Mc4r* gene was reduced accordingly in hypothyroid C57BL/6J mice as it follows the reduction of *Pomc* gene. However, mRNA levels of *Mc4r* remained unchanged in hypothyroid WSB/EiJ mice despite the downregulation of *Pomc* gene. Therefore, the melanocortin pathway seems to be reduced in C57BL/6J mice whereas maintained in WSB/EiJ strain, explaining the weight difference between both hypothyroid strains. Moreover, it has been reported that the expression of *Mc4r* gene is negatively regulated by TH since its expression is upregulated in hypothyroid mice (Decherf *et al*., 2010). This contradicts our results as mRNA level of *Mc4r* gene did not increase in hypothyroid mice. This could be due to the difference in the duration of the treatment that was applied: hypothyroidism was induced for a short period of time (13 days) contrary to our experimental duration (7 weeks). This suggests that negative regulation of *Mc4r* gene by TH may be reversed by prolonged and severe hypothyroidism compromising the activation of anorexigenic mechanisms (as observed in C57BL/6J mice). Another explanation could be that the T3 hypothalamic levels could be maintained by increased D2 activity, which could then counteract the serum hypothyroidism. Consequently, we hypothesized that in WSB/EiJ mice, the melanocortin pathway may be regulated by other mechanisms in response to hypothyroidism allowing mice to maintain their body weight, contrary to C57BL/6J mice. These differential responses revealed between the two strains highlight the resistant phenotype of WSB/EiJ mice to hypothyroidism that relies on their ability to maintain energy homeostasis through adaptive mechanisms.

## Conclusion

In this study, 7 weeks of PTU-treatment induced alteration of lipid metabolism in both strains, particularly a reduction of hepatic lipid synthesis. The metabolic dysregulations induced by hypothyroidism, including this negative energy balance, unlike to what is observed in lipid excess, did not promote peripheral inflammation. However, only WSB/EiJ mice restored their lipid levels by mobilizing lipid machinery in white adipocytes, thus maintaining their body weight. WSB/EiJ mice displayed a phenotype of resistance to metabolic dysregulations induced by hypothyroidism. This compensatorymechanisms, as well as their resistance to HFD-induced obesity (Lee *et al*., 2011; Terrien *et al*., 2019), highlight their adaptive capacities to maintain metabolic homeostasis, namely, their high metabolic flexibility, despite serum hypothyroidism. This model sheds light on the importance of local thyroid homeostasis to maintain lipid metabolism and metabolic homeostasis.

## Materials and methods

### 2.1. Animals and treatment

Animals were housed individually under a 12:12 light dark cycle (07h00-19h00), maintained at 23°C, with food and drinking water provided *ad libitium*. Wild-type C57BL/6J and WSB/EiJ breeder mice were purchased from Charles River (l’Arbresle, France) and Jackson Laboratories (Maine, USA), respectively. Hypothyroidism was induced in 8-weeks old male mice by given an iodine deficient diet supplemented with 0.15% propylthiouracil (PTU, Envigo Teklan, Madison, WI, USA) for 7 weeks as described by (Chaalal *et al*., 2019). These antithyroid molecules block the activity of thyroid peroxidases that catalyse iodination of thyroglobulin and are essential for thyroid hormone synthesis. Moreover, PTU also inhibits deiodinase I (DIO1), which produces T3 by deiodination of T4 in peripheral tissues, such as blood, liver, or kidney (Bianco *et al*., 2002; Cooper, 2005). Euthyroid control (CTRL) mice were fed with standard chow diet. Body and food weights were monitored twice a week during the treatment. All procedures were conducted according to the principles and procedures in Guidelines for Care and Use of Laboratory Animals, and was validated by the MNHN ethical comity for animal experimentations.

### 2.2. Blood and tissue sample collection

At the end of the treatment, retro-orbital blood was collected in the morning, before their euthanasia by decapitation, to measure circulating metabolic parameters. Trunk blood and tissue samples were also collected at this time. Blood samples were centrifuged (15 min, 3 000 x g, room temperature) and serum supernatants were stored at -20°C until analysis. Ependymal white adipose tissue (eWAT) samples were weighted before collection. All tissues were snap frozen in liquid nitrogen and stored at -80°C until further analysis.

### 2.3. Circulating metabolic parameters, thyroid hormone levels and cytokines profile

Serum total T_4_ and leptin concentrations were respectively measured using ELISA kits (Labor Diagnostika Nord (LDN), Nordhorn, Germany; EZML-82K, Millipore Billerica, USA) according to the manufacturer’s instruction. Circulating cytokine and lipid profiles were assayed as described previously (Terrien, Seugnet *et al*., 2019).

### 2.4. Reverse-Transcription qPCR

Total RNA from hypothalamus, liver and eWAT samples (n=6 per group) were extracted using RNABle lysis reagent (Eurobio) and the RNeasy® Mini Kit (Qiagen) according to the manufacturer’s protocol. All RNA samples were quantified using Qubit 2.0 Fluorometer (Invitrogen life technologies) and RNA integrity was evaluated using Agilent 2100 Bioanalyzer. Complementary DNA (cDNA) synthesis was performed using Reverse Transcription Master Mix from Fluidigm® according to the manufacturer’s protocol with random primers. Real time quantitative PCR was carried out with QuantStudio 6 Flex Real-Time PCR System (Applied Biosystems) using TaqMan Universal PCR master mix (Applied Biosystems) and pre-designed TaqMan probes (TaqMan Gene Expression Assays, Applied Biosystems) for all genes listed in **supplementary table 1**. The RT-qPCR reactions for each sample was conducted in duplicates and direct detection of the PCR product was monitored by measuring the increase in fluorescence generated by the TaqMan probe. The qRT-PCR data were analyzed using QuantStudio™ Real-Time PCR Software (version 1.3, Thermo Fisher scientific) and ExpressionSuite software (version 1.0-4, Life technologies). For each tissue, three housekeeping genes were selected based on Vandesompele et al. method (Vandesompele *et al*., 2002) and using SlqPCR package (version 1.42.0). A custom R tool was constructed to measure relative gene expression levels according to the ΔΔCT method (Livak and Schmittgen, 2001) and to perform non-parametric statistical tests (two-way ANOVA with permutation) as described previously (Terrien, Seugnet *et al*., 2019). Graphical representations (boxplot whiskers) were performed using FC values (fold-changes compared to C57BL/6J CTRL) on GraphPad Prism (GraphPad Software Inc., San Diego, CA, USA; version 8.0-2).

### 2.5. Immunohistochemistry

Hypothalamic immunohistochemistry (n=3-4 per group) was assayed as described previously (Terrien, Seugnet *et al*., 2019). Brain sections were observed under TCS-SP5 Leica confocal microscope using x400 magnification. Acquisitions were obtained using max intensity Z projection of 30 μm-thick z-stacks (a minimum of 20 images with a z-step of 1.01 μm) for each hypothalamic arcuate nucleus (ARC) region of interest (ROI; left and right side). Quantifications of microglia and astrocytes were performed with ImageJ software (National Institutes of health, USA) using cell counter plugin. Results of GFAP and IBA1 densities are presented as the number of immuno-positive cells per mm^2^ for each ROI.

### 2.8. Statistical analyses

Statistical analysis for body weight and food consumption were carried out using GraphPad Prism. Variations between strains (C57BL/6J; WSB/EiJ) during the 7 weeks of treatment (CTRL; PTU) were analyzed with the two-way repeated measures ANOVA test followed by

Tukey’s multiple-comparisons test. For the rest of the experiments, non-parametric two-way ANOVA test with permutation was performed using Rscript as described previously (see above). When effect “treatment” and “strain” were only achieved, non-parametric one-way ANOVA test with permutation was applied in order to see in which strain the effect of treatment was driven. Dixon’s Q-test was used for identification and rejection of outliers. Differences were considered significant for a p-value ≤ 0.05. Unless specifically mentioned, all data are represented as medians using boxplot and min-max whiskers on GraphPad.

## Supplementary Figures

**Supplementary Figure 1.**
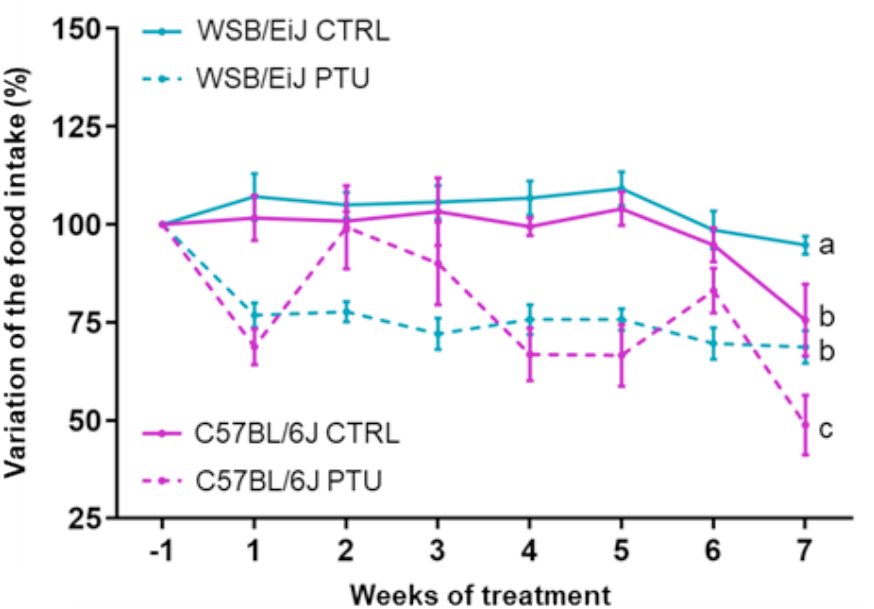
Food intake variation during the treatment. Percentage of the food intake (FI) variation upon weeks of treatment for each group (%FI at each week / Initial FI before the treatment (Week -1); n =7-8 per group). Statistical differences between treatment and time were assessed by two-way repeated measures ANOVA followed by Tukey’s multiple-comparisons test. Significant differences were indicated by different letters to account for group differences at week 7. Values are mean ± SEM.

**Supplementary Figure 2.**
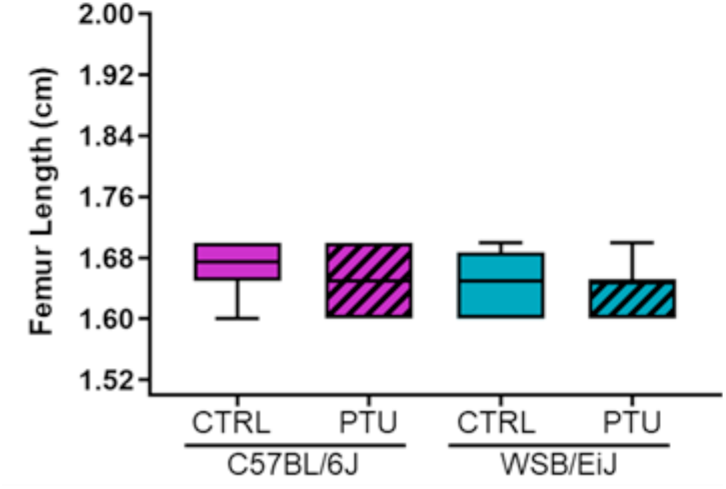
Effect of hypothyroidism on bone growth. Femur length measurements were not affected by PTU-treatment in C57BL/6J and WSB/EiJ mice (n= 8-12 per group; non-parametric two-way ANOVA with permutations test, p > 0.05). Boxplot represents median values and min-max whiskers.

**Table S1:** TaqMan gene expression assays used for qPCR analysis (Related to Figure 2 and Figure 4-7).

| Gene name | Association | TaqMan Gene expression assay ID |
| --- | --- | --- |
| Melanocortin 4 receptor (*Mc4r*) | Central and peripheral metabolic genes (lipid metabolism, food intake, energy metabolism) | Mm00457483_s1 |
| Proopiomelanocortin (*Pomc*) |  | Mm00435874_m1 |
| Agouti related neuropeptide (*Agrp*) |  | Mm00475829_g1 |
| Leptin receptor (*Lepr*) |  | Mm00440181_m1 |
| Peroxisome proliferator activated receptor alpha (*Ppara*) |  | Mm00440939_m1 |
| Peroxisome proliferator activated receptor gamma (*Pparγ*) |  | Mm01184322_m1 |
| Patatin-like phospholipase domain containing 2 (*Pnpla2*) |  | Mm00503040_m1 |
| Peroxisome proliferative activated receptor, gamma, coactivator 1 alpha (*Ppargc1α*) |  | Mm01208835_m1 |
| Fibroblast growth factor 21 (*Fgf21*) |  | Mm00840165_g1 |
| Fatty acid synthase (*Fasn*) |  | Mm00662319_m1 |
| Acetyl-Coenzyme A carboxylase alpha (*Acacα*) |  | Mm01304277_m1 |
| Leptin (*Lep*) |  | Mm00434759_m1 |
| Interleukin 1 beta (*Il1β*) | Inflammatory genes (cytokines, inflammatory pathway) | Mm00434228_m1 |
| Interleukin 10 (*Il10*) |  | Mm00439614_m1 |
| Interleukin 6 (*Il6*) |  | Mm00446190_m1 |
| Tumor necrosis factor (*Tnfa*) |  | Mm00443258_m1 |
| Hypoxanthine guanine phosphoribosyl transferase (*Hprt*) | Housekeeping genes | Mm00446968_m1 |
| Actin beta (*Actb*) |  | Mm00607939_s1 |
| Eukaryotic translation initiation factor 2A (*Eif2a*) |  | Mm01289723_m1 |
| Beta-2-microglobulin (*B2m*) |  | Mm00437762_m1 |
| Phosphoglycerate kinase 1 (*Pgk1*) |  | Mm00435617_m1 |
| Tyrosine 3-monooxygenase/tryptophan 5-monooxygenase activation protein, zeta polypeptide (*Ywhaz*) |  | Mm03950126_s1 |

**Table S2:**
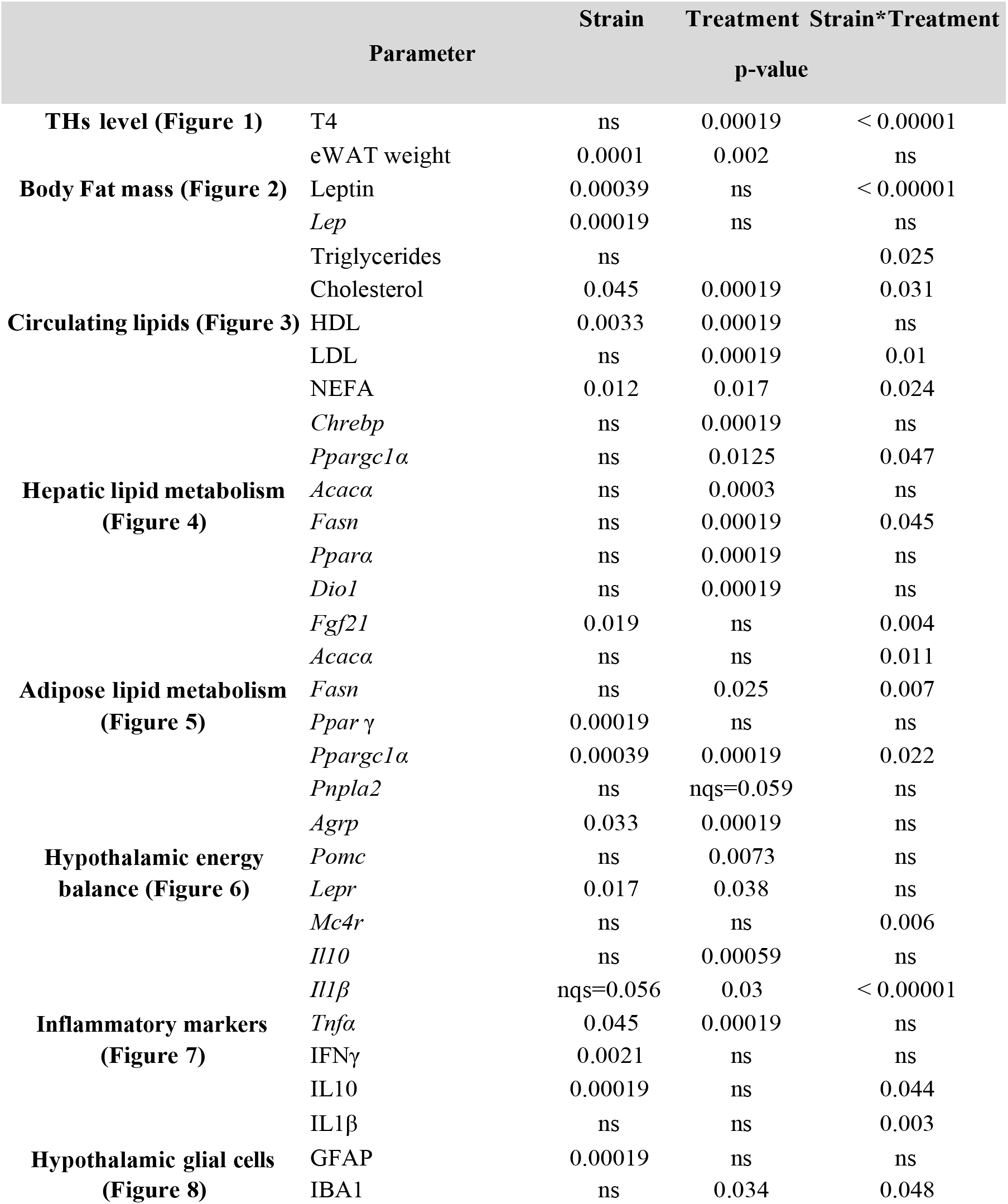
Summary table of the statistics (p-value) for metabolic (related to Figure 1 to Figure 6) and inflammatory parameters (related to Figure 7-8). Statistical non-parametric two-way ANOVA with permutations tests (strain; treatment; Strain*treatment effects) were performed followed by post-hoc analysis (indicated on each graph). nqs= not quite significant; ns= not significant.

## Acknowledgments

Authors wish to thank the valuable technical support from the following French scientifc services: Cytometry and Immunobiology Facility (Cybio, INSERM U1016, Paris) and the Plateforme de Biochimie (CRI, INSERM UMR1149, Paris). We acknowledge the ImagoSeine core facility of the Institut Jacques Monod for the imaging advice. We also thank F. Uridat and S. Sosinski for excellent animal care.

